# Outbreak.info Research Library: A standardized, searchable platform to discover and explore COVID-19 resources

**DOI:** 10.1101/2022.01.20.477133

**Authors:** Ginger Tsueng, Julia L. Mullen, Manar Alkuzweny, Marco Cano, Benjamin Rush, Emily Haag, Outbreak Curators, Jason Lin, Dylan J. Welzel, Xinghua Zhou, Zhongchao Qian, Alaa Abdel Latif, Emory Hufbauer, Mark Zeller, Kristian G. Andersen, Chunlei Wu, Andrew I. Su, Karthik Gangavarapu, Laura D. Hughes

## Abstract

To combat the ongoing COVID-19 pandemic, scientists have been conducting research at breakneck speeds, producing over 52,000 peer-reviewed articles within the first year. To address the challenge in tracking the vast amount of new research located in separate repositories, we developed outbreak.info Research Library, a standardized, searchable interface of COVID-19 and SARS-CoV-2 resources. Unifying metadata from sixteen repositories, we assembled a collection of over 350,000 publications, clinical trials, datasets, protocols, and other resources as of October 2022. We used a rigorous schema to enforce consistency across different sources and resource types and linked related resources. Researchers can quickly search the latest research across data repositories, regardless of resource type or repository location, via a search interface, public API, and R package. Finally, we discuss the challenges inherent in combining metadata from scattered and heterogeneous resources and provide recommendations to streamline this process to aid scientific research.

## Introduction

In early January 2020, SARS-CoV-2 was identified as the virus responsible for a series of pneumonia cases with unknown origin in Wuhan, China^1^. As the virus quickly spread all over the world, the global scientific community began to study the new virus and disease, resulting in the rapid release of research outputs (such as publications, clinical trials, datasets) and resources (*i.e*. research outputs, websites, portals and more). The frequently uncoordinated generation and curation of resources by different types of resource generators (such as government agencies, NGOs, research institutes, etc.) exacerbate four factors that make finding and using resources a challenge: volume, fragmentation, variety, and standardization (**Supplemental Figure 1**). These four factors hamper the ability for researchers to discover these resources, and consequently, impede the translation of these protocols, data, and insights into a synthesized understanding of the virus to help combat the pandemic.

For example, the volume of peer-reviewed articles from a single resource (LitCovid) has grown from about 52,000 published within the first twelve months to over 250,000 as of June 2022. Since April 2020, over 1,000 different research outputs have been published on a weekly basis, spanning new protocols, datasets, clinical trials, as well as publications. The rapid proliferation of resources could be manageable if there were a centralized repository for finding them, but none exists. In addition to research outputs like scientific literature, researchers, public health officials, media outlets, and concerned communities independently developed websites providing highly localized or specialized information on infection rates^2,3^, prevention policies^4,5,6^, and travel restrictions^7^ resulting in a fragmented landscape of very different types of resources (**Supplemental Figure 1**).

The volume and fragmentation issues were immediately obvious. Lacking alternate solutions for addressing these issues, individual and community efforts for curating these resources were created via shared Google spreadsheets^8,9,10^ to aid in discoverability. However, the sheets were not a scalable solution and usually lacked sufficient metadata for describing resources, with the exception of Navarro and Capdarest-Arest^10^. Several projects have attempted to address the volume and fragmentation issues, but were most often focused on a single type of resource. For example, NIH’s iSearch COVID-19 portfolio^11^ and the Kaggle COVID-19 Open Research Dataset Challenge (CORD-19)^12^ aggregate scholarly articles, but do not include clinical trials, datasets, or other types of resources.

Compounding search issues caused by the variety of resource types, there has been a long-standing lack of standardization even *within* a particular type of resource. Existing resource repositories which were able to pivot quickly and curate COVID-19 content from their collections utilized pre-existing metadata standards. For example, researchers involved in PubMed, which uses Medline citation standards, shifted quickly to create LitCovid^13^ which follows the same standard. Similarly, the National Clinical Trials Registry has their own custom list of COVID-19 Clinical Trials which follows their own Protocol Registration and Results System (PRS) schema^14^, but these conventions are not followed by the WHO International Clinical Trials Registry Platform. Zenodo^15^ and Figshare^16^, which both enable export to multiple open data formats including Schema.org, do not completely agree on the marginality, cardinality, and selection of the properties in profiles they use^17,18,19^.

Once the issues of volume, fragmentation, variety, and standardization of resources are addressed, accessibility of the resulting resources for reuse must be addressed. Standardized, centralized resources are of no value if researchers are not able to leverage them. Researchers seeking to process information *en masse* will need an API, while researchers seeking to browse and explore will prefer a user-friendly interface. APIs themselves are less useful without a means of understanding the underlying metadata/data (such as documentation or a GUI), and a user-friendly search portal will be less useful without the inclusion of value-added metadata (such as ones supporting search/filter, linkage and exploration, or qualitative evaluation) for improving resource discovery and interpretation. Interpretability of metadata/data is influenced by the order in which information is presented. To address this challenge, the user interface must encourage exploration which gives users control over the information flow to suit their needs. Lastly, if a user has been able to successfully leverage the standardized, centralized resources, they should be able to easily save and share the results of their efforts.

We address the aforementioned challenges inherent in combining metadata from disparate and heterogeneous resources and making information more interpretable by building outbreak.info, a website which integrates a searchable interface for a diverse, heterogeneous resources which we have collected and standardized (metadata) with surveillance reports on SARS-CoV-2 variants and mutants (data). Our outbreak.info Research Library harvests and harmonizes metadata from sixteen sources encompassing publications, clinical trials, datasets, and more into a single searchable index. Following implementation considerations for FAIRness^20^, our website includes programmatic access via APIs and a standardized metadata interface built off Schema.org. Daily updates ensure that site users have up-to-date information, essential in the midst of a constantly changing research landscape. Our infrastructure is modular, allowing easy addition of new data sources, including based on community contributions. Based on our experience unifying metadata across repositories, we will discuss issues with centralizing, standardizing, and returning resource metadata, epidemiological data, and supporting the use of the metadata/data. In a companion piece, we present our efforts to develop genomic reports to scalably and dynamically track SARS-CoV-2 variants^21^.

## Results

### Standardizing metadata through a schema harmonizing a variety of resource types

We address issues with metadata variety, standardization, and fragmentation by developing a harmonized schema. Schema.org provides a framework to standardize metadata for many different types of data found on the world wide web. However, these standards are not preserved across different types of data. For example, publication providers like PubMed typically use the `‘author’` property in their metadata, while dataset providers like Figshare and Zenodo are compliant with the DataCite schema and typically prefer `‘creator’`. Although both properties are valid for their respective Schema.org classes, we normalized our schema to use `‘author’` for all six of our classes (Dataset, ClinicalTrial, Analysis, Protocol, Publication, ComputationalTool) since we expected the volume of publications to dwarf all other classes of resources. We developed a schema that encompassed six types of resources based on their proliferation at the beginning of the pandemic and their importance to the research community: Publications, Datasets, Clinical Trials, Analysis, Protocols, and later ComputationalTools. We added this schema to the Schema Registry of the Data Discovery Engine (DDE)^22^, a project to share and reuse schemas and register datasets according to a particular schema. Using this schema we ingested and harmonized metadata from an initial set of sixteen key resources: LitCovid (Publications), bioRxiv and medRxiv (Publications), COVID-19 Literature Surveillance Team (COVID-19 LST) (Publications), ClinicalTrials.gov (NCT) (ClinicalTrials), WHO International Clinical Trials Registry Platform (WHO ICTRP) (ClinicalTrials), Figshare (Datasets, Publications, and more), Zenodo (Datasets, ComputationalTools, Publications, and more), MRC Centre for Global Infectious Disease Analysis (Analyses, Publications, and More), Protocols.io (Protocols), Protein Data Bank (PDB) (Datasets), Data Discovery Engine (DDE) (Datasets), Harvard Dataverse (Datasets), ImmPort (Datasets), bio.tools (ComputationalTools), and Dockstore (ComputationalTools) (**Figure 1a**).

**Figure 1.**
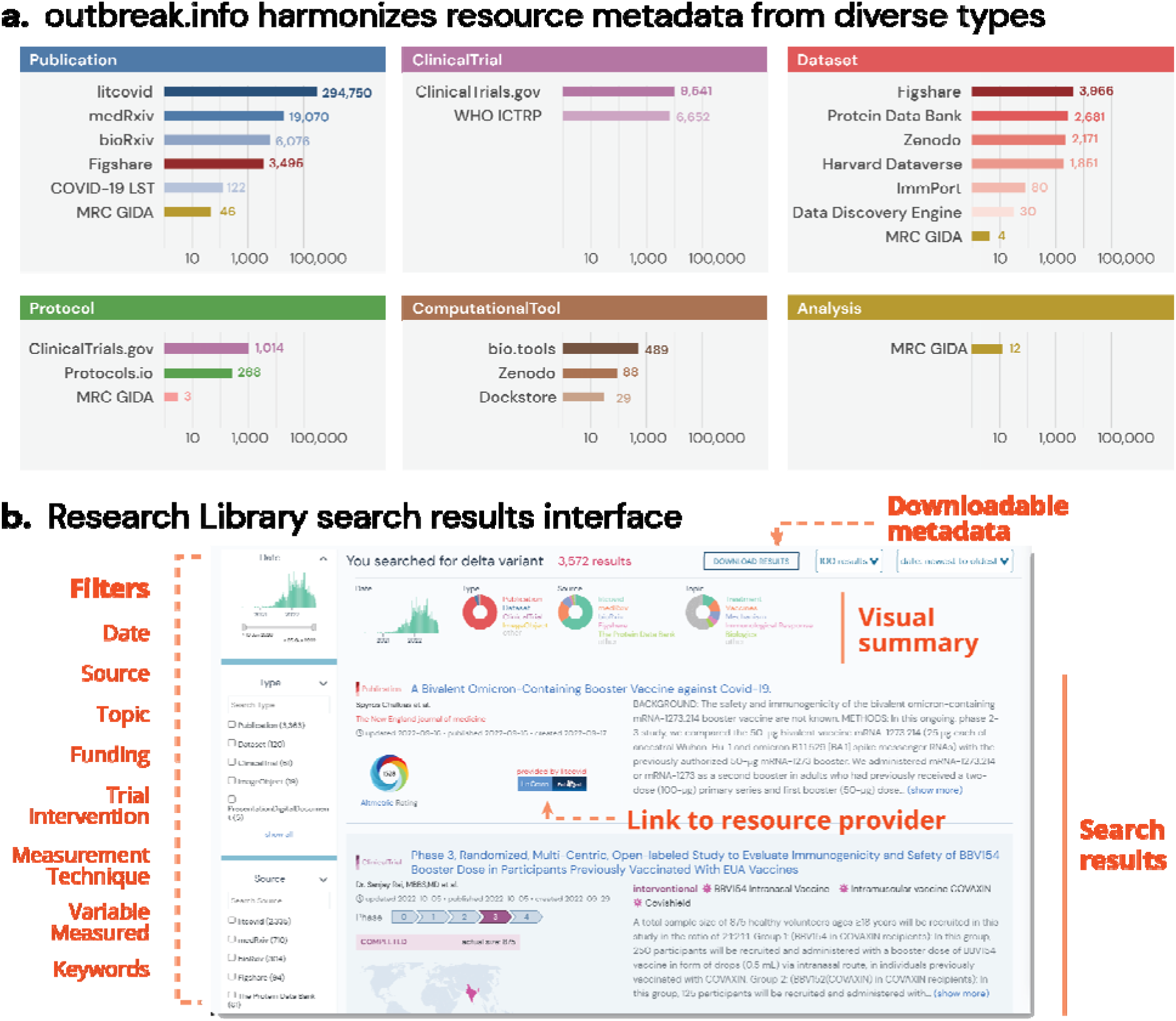
Supporting resource centralization and standardization by developing a harmonizing schem**a. a**, Distribution of resources by resource type and source. **b**, Heterogeneous and filterable resources (i.e. publications, clinical trials, datasets, etc.) resulting from a single search of the phrase “Delta Variant”.

Sources of certain metadata did not map readily to existing Schema.org classes. For example, clinical trials registries like NCT have one general schema for both observational and interventional studies, while Schema.org provides separate classes for each of these types of studies. Since NCT was a primary source of clinical trials metadata for our research library, we tailored the Outbreak schema based on the combined general NCT schema. Fortunately, many dataset repositories (including Harvard Dataverse, Figshare, and Zenodo) offered Schema.org-compliant metadata, even if the repositories differed in the metadata fields that were available. Once our schema was developed, we created parsers (data plugins) to import and standardize metadata from our initial set of resources. We assembled the data plugins into a single API via BioThings SDK^23^, and scheduled them to update on a daily basis to ensure up-to-date information. All *structured* metadata provided by the data source was saved and harmonized in this protocol, and the completeness of each metadata property by resource type is shown in **Supplemental Figure 2** (data provided in **Supplemental Table 1**).

By leveraging the BioThings SDK, we developed a technology stack that addresses the fragmentation issue by easily integrating metadata from different pre-existing resources. With a unified schema that harmonizes information across heterogeneous resource types, a single search (for example “Delta variant”) to our API can return relevant publications, datasets, clinical trials, and more (**Figure 1b**). This allows us to create visualizations which allow users to quickly familiarize themselves with the search results. For example, the histogram in Figure 1b indicates that the number of resources mentioning the “Delta variant” began growing mid 2021, and started declining in the summer of 2022. The donut charts indicate that the majority of resources on the “Delta variant” are publications coming from LitCovid.

Although other resources which aggregate heterogeneous types have since been developed, they do not offer the same types or features as our site, including filtering, sorting, visual summaries, downloadable metadata, and an API for programmatic access, and rarely cover the breadth of research types supported in the Research Library (**Supplemental Table 2**). A comparison of features from our Research Library with other COVID-19 multisource aggregation efforts can be found in **Supplemental Table 2**, and a A list of terms and features are defined in **Supplemental Table 3**. As seen in **Supplemental Table 4A**, the most common searched source (*i.e*.-filter by source) has been LitCovid (website) and bioRxiv (API); while the most common resource types searched (*i.e*.-filter by resource type) has been for publications, datasets, and clinical trials. Usage stats for record views and filtering by source are available in **Supplemental Table 4B**. Filtering was the most popular feature added to the Library, with over a quarter of all queries using some sort of filtering (**Supplemental Table 4C**). Users were most likely to filter results by resource type, followed by keywords and source.

### Enabling community curation and metadata submission to address fragmentation and standardization issues

At the start of the pandemic many curation efforts were neither coordinated, standardized, or easy to find; however, these efforts served an important role in organizing information early on. Given the highly-fragmented, diffuse and frequently changing nature inherent to biomedical resources, we built outbreak.info with the idea that it should be expanded with the participation of the community. Not only is finding and adding resources to the collection an onerous process, it also requires us to know the full landscape of resources on the internet. Furthermore, many resources do not collect metadata useful for linkage, exploration, and evaluation in machine-readable formats. We enabled community-based contributions of resource metadata in a variety of ways (**Figure 2a**).

**Figure 2.**
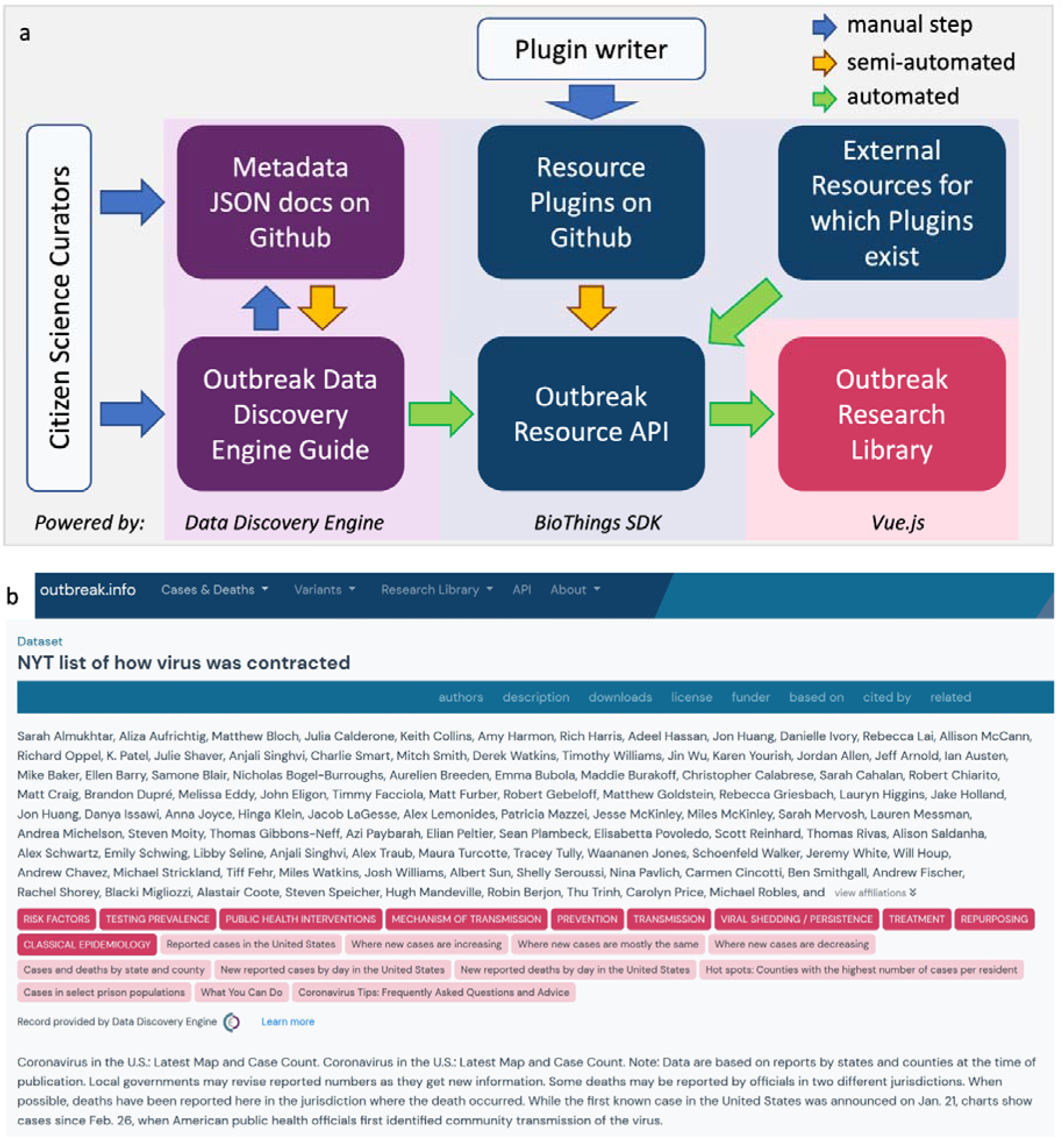
Aggregating resource metadata by leveraging community contributions. **a**, The community contribution pipeline and technology stack for outbreak.info’s Research Library. Curators may submit dataset metadata using the DDE built-in guide or from GitHub via the DDE/BioThings SDK. Python-savvy contributors can create parsers to contribute even more metadata via the BioThings SDK plugin architecture. A resource plugin allows the site to automatically ingest and update metadata from the corresponding external resource. Blue arrows indicate manual steps, yellow arrows indicate automatable steps after an initial set up, green arrows indicate completely automated steps. **b**, An example of a detailed metadata record manually-curated by volunteers as it appears in the Research Library.

For single datasets, contributors can submit the metadata via outbreak.info’s dataset submission guide on the Data Discovery Engine^22^, which ensures that the curated metadata conforms to our schema. From there, it can be saved to GitHub, where it can be improved by other contributors via forking and pull requests. The DDE automatically passes the information to the outbreak.info Resources API where it is made discoverable with the Research Library. We demonstrated its utility by asking two volunteers to annotate metadata from thirty different individual resources from across the internet and submitted the metadata for integration via the DDE. As seen in **Figure 2b**, community-contributed metadata using the DDE is standardized and can be exhaustively detailed. Although both of our volunteers provided values for many of the available metadata properties (`name, description, topiccategories, keywords`, etc.), one provided an extensive list of authors. Using the BioThings SDK in conjunction with the DDE allows us to centralize and leverage individualized curation efforts that often occur at the start of a pandemic. Additionally, collections of standardized datasets, publications and other resources can be submitted to the Outbreak Resources API by contributing a resource plugin. Resource plugins are BioThings-compatible Python scripts to harvest metadata from a source and standardize it to our schema; these parsers can be submitted by anyone with Python coding skills^23^. Our community contribution pipeline allows us to quickly and flexibly integrate the uncoordinated data curation efforts, particularly apparent at the start of the pandemic (**Supplemental Figure 3**).

### Improving searching, linkage and evaluation of resources to support exploration

Centralizing and standardizing the resources does not automatically make the resources explorable to a user. While centralizing and standardizing allows for search, aggregation and some filtering; additional metadata and a user-friendly interface is needed to allow thematic browsing/filtering and to enable iterative traversal from query to search result to refined query and vice versa. To ensure ease of access to and ease of use of our research library, we conducted usability studies and iteratively improved our site (**Supplemental Figure 4**). To support resource exploration and interpretation, we added properties (value-added metadata) to every class in our schema that would support searching/filtering/browsing (`topiccategories`), linkage/exploration (`correction, citedBy, isBasedOn, isRelatedTo`), and interpretation (qualitative evaluations) of resources.

We selected these properties based on pre-existing citizen science and resource curation activities, suggesting their value in promoting discoverability. For example, citizen scientists categorized resources in their lists/collections by type (Dataset, ClinicalTrials, etc.) in their outputs^10^ or area of research (Epidemiological, Prevention, etc.)^24^ as they found these classifications helpful for searching, filtering, and browsing their lists/collections. They also evaluated the level of evidence provided by these resources in order to improve its interpretability (*i.e*. understanding the credibility/quality of the resource)^24^. Existing repositories such as LitCovid also organized information to enhance browsability, but these efforts were often not captured in the metadata. For instance, LitCovid organized publications into eight research areas such as Treatments or Prevention, but these classifications are not available in the actual metadata records for each publication. To obtain these classifications from LitCovid, subsetted exports of identifiers must be downloaded from LitCovid and then mapped to the metadata records from PubMed.

To classify resources by `topiccategory` and improve search/browse/filtering capabilities in our user interface, we used a combination of existing work (LitCovid) and human curation to augment that categorization to provide higher specificity of topics and to extend to new types of data (datasets, clinical trials). We applied out-of-the-box logistic regression, multinomial naive bayes, and random forest algorithms from scikitlearn to classify each resource as belonging or not belonging to each topic. A resource was only classified as belonging to each topic if all three algorithms agreed on the classification. These three algorithms were found to perform best on this binary classification task using out-of-the-box tests. For example, if a user wants to browse for all resources (or filter down search results) related to the prevention of COVID-19, they can select the appropriate `topiccategory` in the search/search results view of the resources (**Figure 3a**). Users can also easily traverse from a view of a resource record to start a new search by clicking on a `topiccategory` of interest (**Figure 3b**). We further enable exploration by populating the linkage properties (`corrections, citedBy, isBasedOn, isRelatedTo`) from citation metadata (whenever possible), corrections metadata (from LitCovid, when available), and via an algorithm for matching peer-reviewed papers in LitCovid with their corresponding preprints in bioRxiv/medRxiv. Together with the corrections metadata from LitCovid, the algorithm has matched over 2,600 peer-reviewed articles with their corresponding preprints, enabling users to follow from a publication record from LitCovid to a publication record in bioRxiv/MedRxiv (**Figure 3b**).

**Figure 3.**
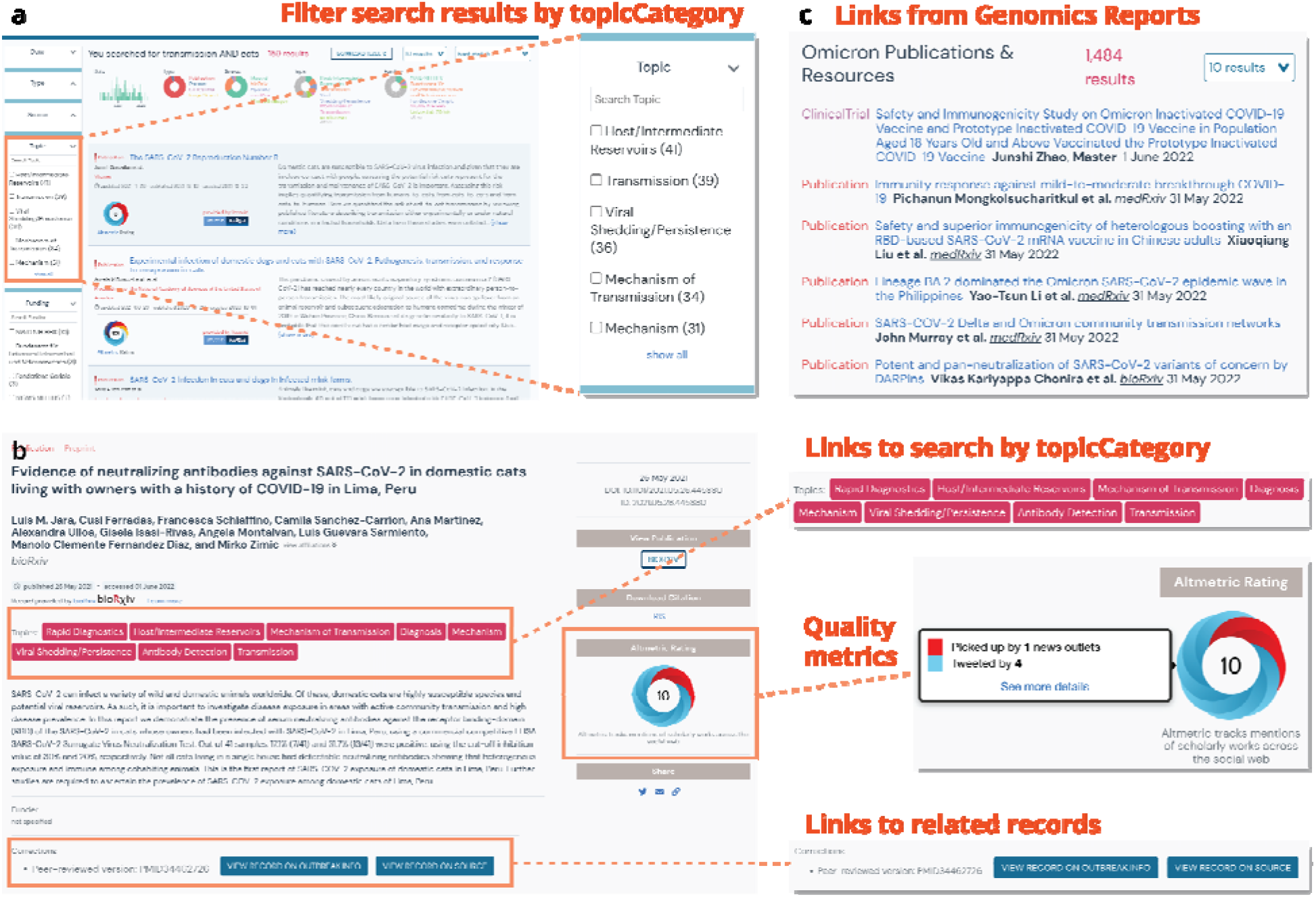
Enabling exploration of the resources. **a**, Selectable options for filtering results by topic category or other facets enhance searchability and exploration from the search results view. **b**, Links to other records or to additional potential searches of interest enabling further exploration from a record view. **c**, Links from the Omicron Variant report to related resources.

Once a user has found a record of interest, they might wonder about the credibility of the resource. To populate resource evaluations so that users can assess the quality of a resource and tailor their interpretation accordingly, we leveraged the Oxford 2011 Levels of Evidence annotations generated by the COVID-19 Literature Surveillance (COVID-19 LST) team^24^ as well as Digital Science’s Altmetrics^25^. These evaluations are currently visible in the search results, and in the future, we will enable users to further filter or sort search results by some measurement of quality (*i.e*. Altmetrics: degree of access, or COVID-19 LST: level of evidence). Lastly, we integrated resources with data and analyses we curated to track SARS-CoV-2 variants^21^. Researchers can seamlessly traverse from a specific variant report like Omicron to resources on that variant to help understand its behavior (**Figure 3c**). In the absence of a centralized search interface with linked records, a similar attempt to explore resources outside the outbreak.info portal would require extensive manual searching from multiple different sites (**Supplemental Figure 5**), each with their own interfaces and corresponding search capabilities.

### Case Study: Research on SARS-CoV-2 variants

To demonstrate the unique features of the outbreak.info Research Library, we explored the dynamics of research into SARS-CoV-2 variants over time. In particular, we focus on the ability to rapidly identify new research on an evolving topic; the benefits of querying across repositories and types using a standardized interface; the integration of quality metrics to evaluate results; and the ability to investigate hypotheses about research trends using our API via the R package.

As SARS-CoV-2 continues to evolve, Variants of Concern (VOCs) have emerged with increased transmissibility, virulence, and/or immune evasion. These variants have led to worldwide waves of cases and deaths and an expanding interest within the research community to understand their behavior; searches on particular mutations, lineages, and VOCs remain some of the most commonly searched terms in the Research Library (**Supplemental Table 4**). Using the Research Library, we sought to answer two key questions: (***1***) How has the research community responded to the emergence of new variants? (***2***) How has that response changed over time?

We extracted research related to variants in the Research Library using the query variant OR lineage. Since our library combines metadata across 16 sources, we can simultaneously query across each of these sources, and as well as across different research types (**Figure 4a**). Over 10,000 separate entries about variants are within the Library as of October 2022, including Publications, Datasets, Clinical Trials, Protocols, and more. Using filters and the quality metrics provided through Altmetric badges, we can quickly identify which results have been recognized by the community via Altmetric scores, such as an RT-qPCR protocol to screen VOCs (**Figure 4b**). Clearly, variants are an active area of research; the question is, has this enthusiasm changed over time? Using the outbreak.info R package, we can access the harmonized metadata assembled in the Research Library to execute more complex analyses. By examining the proportion of research related to variants in the Research Library over time, we find that there was an increase in research on variants following the first identification VOCs such as Alpha (B.1.1.7*) and Beta (B.1.351*) (**Figure 4c**). This increase was even more prominent for the Omicron (B.1.1.529*) variant in late 2021; we hypothesize that this increase was due to the heightened awareness of the value in studying variants amongst the scientific community, and early indications that the variant could be of global concern (high growth rate of Omicron and the presence of a large number of mutations in important sites). To examine how research differed by VOC over time, we constructed queries for each VOC, including its Pango lineage name and associated sublineages. To ensure our search results focused on the variant and to avoid results that just mentioned the term in their description, we restricted our queries to only the name field. With the three VOCs which became the dominant worldwide form of SARS-CoV-2 (Alpha, Delta, and Omicron), we find that the increase in research on these VOCs mirrors the rise in worldwide prevalence for each variant, with the research output roughly proportional to global prevalence (**Figure 4d**). With Alpha and Delta, there was a slight lag in research publications which was not observed with Omicron, and research on Omicron over the last ten months has dwarfed the other VOCs. Lastly, research on previously circulating variants (Alpha, Beta, Gamma, Delta) continues, even though these variants are rarely detected presently. Research continues on these near-extinct VOCs in four areas, complementary to ongoing research on Omicron: retrospective analyses, fundamental studies on the mechanisms of action, comparative studies related to the currently circulating variant Omicron, and studies of recombinant variants. Taken together, the research community’s response to the emergence of new variants has been robust, has become a greater focus of overall research effort, and quickly pivots to studying the dominant variant.

**Figure 4.**
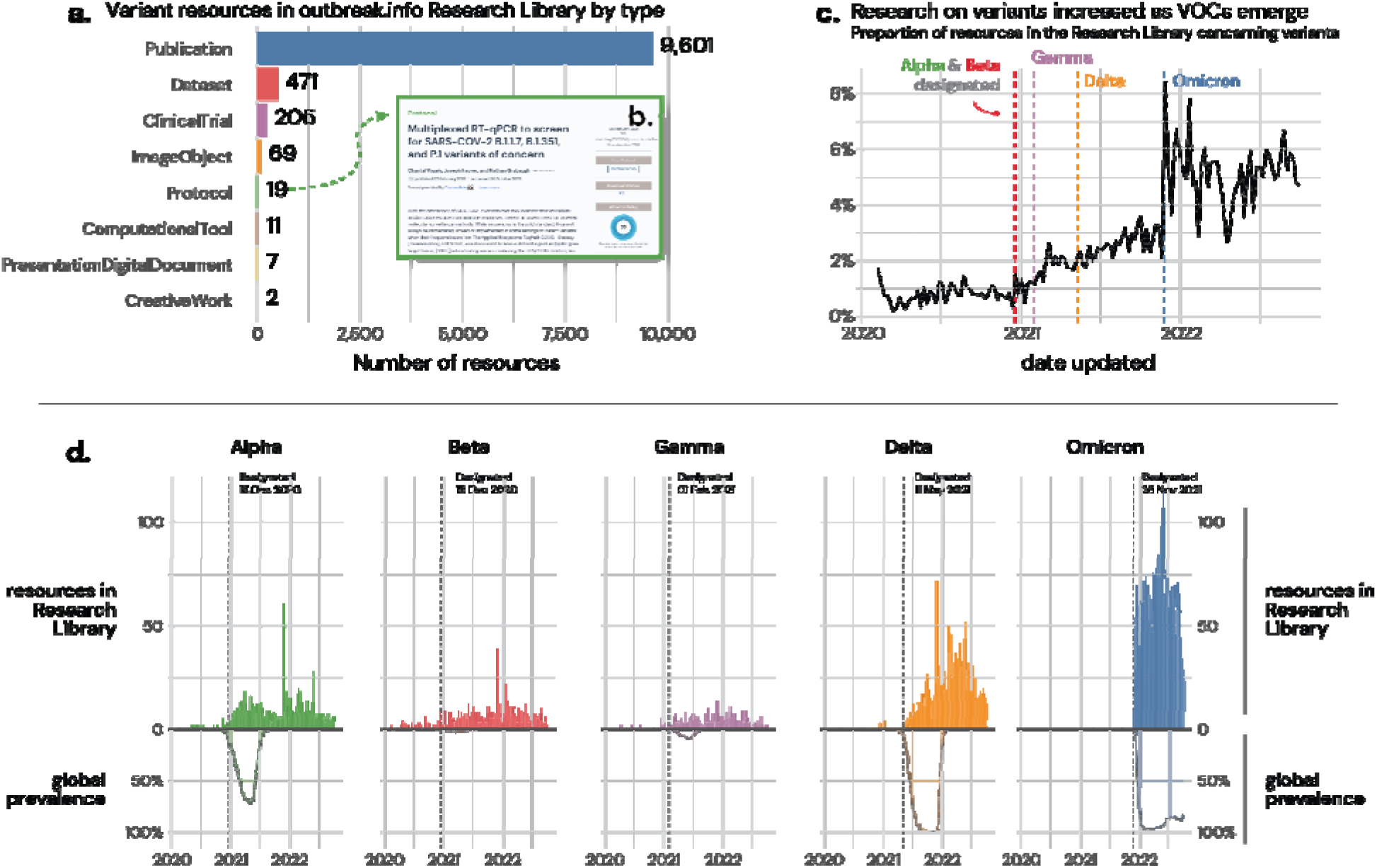
Resources concerning variants within the outbreak.info Research Library. **a**, The Research Library harvests metadata from 16 sources, enabling standardized searching across resource types, including Publications, Datasets, ClinicalTrials, Protocols, and more. **b**, An example variant protocol discovered within the Library. **c**, As Variants of Concern (VOCs) were designated, the proportion of research in the Library focused on variants increased. **d**, The increase in research on each VOC mirrored its worldwide prevalence, with research on the transmissibility, virulence, and/or immune evasion supporting their VOC designation by public health agencies, and these designations encouraging further research.

## Discussion

Over the course of the COVID-19 outbreak, researchers have shared the results of their work at unprecedented levels – exacerbating existing issues in resource volume, fragmentation, variety, and standardization. These issues make it challenging to assemble, traverse, and maintain up-to-date resources. Further, the urgency of a pandemic requires that these issues be addressed quickly, and in a scalable manner to be able to accommodate more data flexibly. We launched outbreak.info within two months of the start of the COVID-19 pandemic to address these issues and to highlight barriers in rapidly sharing research outputs in the midst of a pandemic.

To address the structure and standardization issue, we developed a standardized schema, integrated all structured metadata from different resources into an openly accessible API, and created a user-friendly search-and-filter, web-based interface. In addition to difficulties standardizing inconsistent metadata models between resources, it is also challenging to maintain a resource library that imports metadata from so many sources, particularly when the metadata updates daily and is prone to frequent changes in the structure of the data. Any changes to the upstream metadata offered by an external site necessitates a change in the parser which imports them. The resource API infrastructure (**Supplemental Figure 6**) utilizes the BioThings SDK plugin architecture to handle errors in individual parsers without affecting the availability of the API itself. Errors thrown by individual parsers may result in a lack of updates of an individual resource until the error is resolved, but the API will serve the latest version of data from the broken parser and up-to-date data from all functional parsers, which will continue to be updated independently. Using the plugin architecture also allows the creation and maintenance of the individual resource parsers to be crowdsourced to anyone with basic Python knowledge and a GitHub account. Although resource plugins allow outbreak.info to ingest large amounts of standardized metadata, there are still many individual datasets and research outputs scattered throughout the web which are not located in large repositories. Since it is not feasible for one team to locate, identify, and collect standardized metadata from these individual datasets and research outputs, we leveraged the Data Discovery Engine^22^ to enable crowdsourcing and citizen science participation in the curation of individual resource metadata.

At the onset of our data harvesting and harmonization efforts, we focused on creating a unified search interface backed by a common Schema.org-based schema. With an extendable pipeline to harvest metadata and an interface to search them in place, we focused next on augmenting the existing metadata by adding properties to help researchers find information more quickly: topic categorization to group related research, resource linking to connect preprints to their published articles, and integrating external evaluations of the research trustworthiness using a combination of human curation and automated methods.

Citizen scientists have played an active role in data collection^26,27^ and making information more accessible^12,24^ throughout the ongoing pandemic. Given their ability to perform information extraction^28^ and their immense contributions to classification tasks^29^, we incorporated citizen science contributions into the training data for classifying resources into our topic categories. Some resource aggregators have used clustering algorithms to categorize the entries in their resource libraries, though many only aggregate resources of a single type (i.e. publications). We employed an approach based on citizen scientist classification due to the heterogeneity of our resources which leverages previous classification work from LitCovid. Publications in LitCovid are already categorized into LitCovid categories which were readily used as training data for our topic category classifier. Further, LitCovid publications are indexed in PubMed and often include MeSH terms which could be mapped and used as training data. In contrast, clinical trial records lacked such explicit classifications, but may have metadata regarding study types, design, arm groups, and interventions which can be mapped to topic categories and used as training data. Datasets shared early on in the pandemic often provided minimal metadata, which is why we turned to citizen science classifications to address the heterogeneity in metadata coverage. Our API ensures that all records are openly accessible for downstream analyses, so anyone is welcome to apply their preferred approach to organizing the data.

In addition to generating metadata values for improved searching and filtering, we enabled linkages between resources in our schema. For instance, ideally a publication about a clinical trial would link to its clinical trial record, protocols used to collect the data, datasets used in their analyses, and software code underlying the analyses to enable a more meaningful understanding of this trial. However, these connections rarely exist within the metadata; as a result, we have generated linkages between preprints and peer-reviewed publications, and plan to create more linkages between other resource types. Challenges to include these linkages included: the lack of unique identifiers, inconsistent use of citation metadata fields between resources, and the lack of structured linkage metadata. For example, the ONS Deaths Analysis does not have a unique identifier as assigned by Imperial College London, lacks any citation metadata fields, and instead mentions a potential linkage to an Imperial College London report in its mention of limitations^30^. Although preprints from bioRxiv^31^ and medRxiv may have links to the corresponding peer-reviewed manuscript on the bioRxiv site, this information is not accessible via their API, necessitating the use of algorithms to generate these links.

As a result of this centralization, standardization, and linkage, the outbreak.info Research Library and resources API has been widely used by the external community, including journalists, members of the medical and public health communities, students, and biomedical researchers^32^. For instance, the Radx-Rad Data Coordination Center (https://www.radxrad.org) is utilizing the Outbreak API to collect articles for customized research digests for its partners. Using the Radx-Rad SearchOutbreak app (https://searchoutbreak.netlify.app), users select topics based on information submitted from partners. These are turned into queries for the Outbreak API, and every week, new articles are added to the digests which are available at the website. A workflow sends an email to subscribed users. These digests are not currently available to the public but are expected to be released publicly in the future^33^. On average, the Research Library receives nearly 3,000 pageviews per month, of which 85% are unique visitors (**Supplemental Table 4**). The Research Library site has been used for over 11,000 unique searches and the Research Library API receives an average of nearly 63,000 unique hits per month (including web traffic and programmatic access). As with many web-based, open source resource sites, bugs and browser-compatibility issues may arise without notice for less popular browsers. Users can bring these issues to our attention by submitting them to our issue tracker on GitHub (https://github.com/outbreak-info/outbreak.info/issues).

While we have developed a framework for addressing resource volume, fragmentation, and variety that can be applicable to future pandemics, our efforts during this framework exposed additional limitations in how data and metadata are currently collected and shared. Researchers have embraced preprints, but resources (especially datasets and computational tools) needed to replicate and extend research results are not linked in ways that are discoverable. Although many journals and funders have embraced dataset and source code submission requirements, the result is that the publication of datasets and software code are still heavily based in publications instead of in community repositories with well-described metadata to promote discoverability and reuse. In the outbreak.info Research Library, the largest research output by far is publications, while dataset submission lags in standardized repositories encouraged by the NIH such as ImmPort, Figshare, and Zenodo. We hypothesize that this disparity between pre-print and data sharing reflects the existing incentive structure, where researchers are rewarded for writing papers and less for providing good, reusable datasets. Ongoing efforts to improve metadata standardization and encourage schema adoption (such as the efforts in the Bioschemas community) will help make resources more discoverable in the future – provided researchers adopt and use them. For this uptake to happen, fundamental changes in the incentive structure for sharing research outputs may be necessary.

As of September 2022 (33 months since SARS-CoV-2 was first identified as the infectious agent of the COVID-19 pandemic), there have been over 580 million cases and over 6.4 million deaths. As those numbers continue to grow, so too does the research and understanding of the causes and consequences of the spread of this virus – much of which is shared publicly, in near real time. While the unprecedented amount of research on COVID-19 offers new opportunities to accelerate the pace of research, the difficulty in finding research amidst this “infodemic” remains a fundamental challenge. In the outbreak.info Research Library, we address many of these challenges to assemble a collection of heterogeneous research outputs and data from distributed data sources into a searchable platform. To query this unified source of COVID-19 research, we built a web-based search interface to perform cross-repository and cross-type queries to simultaneously search for publications, clinical trials, datasets, computational tools, protocols, and analyses from 16 sources. By making these metadata more finable, accessible, interoperable, and reusable (FAIR), we increase the accessibility of COVID-19 research and enable researchers to quickly find information – essential in the midst of a rapidly changing pandemic. Our metadata processing platform is modular, allowing easy extension to add new metadata sources, allowing the Research Library to grow with the pandemic as research changes. To enable further analysis, we enable programmatic access to the standardized library. For example, we explored trends in research on variants, demonstrating that the research community has pivoted to studying variants after the discovery of the Alpha variant, and especially after the emergence of the Omicron variants. Lastly, with the embrace of open science stored in decentralized sources, quickly finding information will be critical for the *next* pandemic. Our approach to unify metadata across repositories will serve as a template for rapidly creating a unified search interface to aggregate research outputs for any pathogen or any research domain.

## Methods

### Schema development

The development of the schema for standardizing our collection of resources is as previously described^22^. Briefly, we prioritized six classes of resources which had seen a rapid expansion at the start of the pandemic due to their importance to the research community: Publications, Datasets, Clinical Trials, Analysis, Protocols, and Computational Tools. We identified the most closely related classes from Schema.org and mapped their properties to available metadata from 2-5 of the most prolific sources. Additionally, we identified subclasses which were needed to support the aforementioned six classes and standardized the properties within each class. In addition to standardizing ready-to-harvest metadata, we created new properties which would support the linkage, exploration, and evaluation of our resources. Our schema was then refined as we iterated through the available metadata when assembling COVID-19 resources. The Outbreak schema is available at https://discovery.biothings.io/view/outbreak.

### Assembly of COVID-19 resources

The resource metadata pipeline for outbreak.info includes two ways to ingest metadata (**Supplemental Figure 6**). First, metadata can be ingested from other resource repositories or collections using the BioThings SDK^23^ data plugins. For each resource repository/collection, a parser/data plugin enables automated import and updates from that resource. To import the data, the metadata is harvested from the source using API calls (if available), HTML web scraping, or .csv or .txt tables of metadata. All structured metadata provided by the sources is compiled and mapped to our schema using custom Python scripts. The harmonized metadata is dumped into a JSON output. **Supplemental Figure 2** shows the completeness of each metadata property within our schema, broken down by resource type. Data plugin code for the sources is available at https://github.com/outbreak-info.

In the second mechanism, metadata for individually curated resources can be submitted via an online form through the Data Discovery Engine (DDE) Metadata Registry^22^. To assemble the outbreak.info collection of resources, we collected a list of over a hundred separate resources on COVID-19 and SARS-CoV-2. This list (**Supplemental Table 5**) included generalist open data repositories, biomedical-specific data projects including those recommended by the NIH^34^ and NSF^36^ to house open data, and individual websites we came across through search engines and other COVID-19 publications. Prioritizing those resources which had a large number of resources related to COVID-19, we selected an initial set of 2-3 sources per resource type to import into our collection. Given the lack of widespread repositories for Analysis Resources, only one source would be included in our initial import (Imperial College London). An Analysis resource is defined as a frequently-updated, web-based, data visualization, interpretation, and/or analysis resource.

### Creation of the Research Library API and query interface

In order to accommodate a large number of heterogeneous data sources, each of which is independently harvested, we used the BioThings SDK framework to combine the data sources into a combined searchable index (**Supplemental Figure 6**). The JSON outputs of our data plugins are ingested by the BioThings framework, merged into an intermediary MongoDB database, and the processed data is indexed in an Elasticsearch index. A Tornado server is used to create an API endpoint, api.outbreak.info/resources, that leverages the search capabilities of Elasticsearch to efficiently query data. Within the search results, Elasticsearch sorts them by relevance based on Lucene’s Practical Scoring Function37, which prioritizes the query normalization factor, coordination factor, term frequency, inverse document frequency, and any custom query boosting fields selected by the user. To adjust this behavior based on common search patterns, we upweighted queries where the search term occurs in the name and/or intervention name fields (`weight of name: 4 and interventions.name: 3`). We continue to monitor common query patterns using our analytics to refine the scoring algorithm to improve the list of results for the user. Within the web interface, the user has the option to sort by the best match relevance score, update date for the document, or alphabetically by name. Within search queries, terms are automatically combined by AND; for instance, the search `long covid` will be interpreted as `long AND covid`. Terms can be explicitly combined by the term OR, and exact phrases can be encapsulated in quotes (*e.g*. (`Moderna OR Pfizer`) AND (`“side effects” OR “adverse effects”`)). Further details on advanced searching behavior is provided in our guide to the outbreak.info R package at https://outbreak-info.github.io/R-outbreak-info/articles/researchlibrary.html#some-notes-on-constructing-queries.

To update the API with new data provided by the data sources, the BioThings Hub schedules daily updates to pull data upstream and add them to the existing index. The BioThings Hub independently maintains each data source, enabling independence if an individual data source pipeline breaks, and maintains historical data by default, creating automated backups. The code for the server-side application is available at https://github.com/outbreak-info/outbreak.api.

### outbreak.info Research Library web application and metadata access

The web application was built using Vue.js, a model–view–viewmodel JavaScript framework which enables the two-way binding of user interface elements and the underlying data allowing the user interface to reflect any changes in underlying data and vice versa. The client-side application uses the high performance API to interactively perform operations on the database. To iteratively improve the interface, we conducted usability studies, as described in **Supplemental Figure 4**.The code for the client-side application is available at: https://github.com/outbreak-info/outbreak.info. To enable programmatic access to all our harmonized metadata collection, all data is available in our api, api.outbreak.info, and can be accessed through an R package as described in Gangavarapu *et al*. (package website: https://outbreak-info.github.io/R-outbreak-info/; code: https://github.com/outbreak-info/R-outbreak-info).

### Community curation of resource metadata

Resource plugins such as those used in the assembly of COVID-19 resources do not necessarily have to be built by our own team. We used the BioThings SDK^23^ and the Data Discovery Engine^22^ so that individual resource collections can be added by writing BioThings plugins that conform to our schema. Expanding available classes of resources can be done easily by extending other Schema.org classes via the DDE Schema Playground at https://discovery.biothings.io/schema-playground. Community contributions of resource plugins can be done via GitHub. In addition to contributing resource plugins for collections/repositories of metadata, users can enter metadata for individual resources via the automatic guides created by the Data Discovery Engine. To investigate potential areas of community contribution, we asked two volunteers to inspect 30 individual datasets sprinkled around the web and collect the metadata for these datasets. We compared the results between the two volunteers and their combined results were subsequently submitted into the collection via the Data Discovery Engine’s Outbreak Data Portal Guide at https://discovery.biothings.io/guide/outbreak/dataset. Though limited by the original submission form (Google forms), the raw and merged responses illustrating the thoroughness of the of the submissions from the two volunteers can be found at: https://docs.google.com/spreadsheets/d/1q1c400UFIOyXedFf2L81zROVkXi3BWBhU46Ic0cMYsI/edit?usp=sharing. Improvements or updates for manually curated metadata can be submitted via GitHub pull requests.

### Community curation of searching, linkage, and evaluation metadata and scaling with machine learning

In an effort to enable improved searching and filtering, we developed a nested list of thematic or topic-based categories based on an initial list developed by LitCovid^13^ with input from the infectious disease research community and volunteer curators. The list consists of 11 broad categories and 24 specific child categories. Whenever possible, sources with thematic categories were mapped to our list of categories in order to develop a training set for basic binary (in group/out group) classifications of required metadata fields such as (title, abstract and/or description). If an already-curated training set could not be found for a broad category, it would be created via an iterative process involving term/phrase searching on LitCovid, evaluating the specificity of the results, identifying new search terms by keyword frequency, and repeating the process. To generate training data for classifying resources into specific topic categories, the results from several approaches were combined. These approaches include direct mapping from LitCovid research areas, keyword mapping from LitCovid, logical mapping from NCT Clinical Trials metadata, the aforementioned terms search iteration, and citizen science curation of Zenodo and Figshare datasets. Details on the logical mapping from NCT Clinical Trials metadata can be found at https://github.com/gtsueng/outbreak_CT_classifier. The keyword mapping from LitCovid can be found at https://github.com/outbreak-info/topic_classifier/tree/main/data/keyword and https://github.com/outbreak-info/topic_classifier/tree/main/data/subtopics/keywords. While positive categorical data were identified via the aforementioned methods, negative controls were generated by randomly selecting from alternative topics and ensuring no overlap. The categorical data were randomly split into training (80%) and test (20%) sets per test, and five tests were performed per topic using three methods (logistic regression, multinomial naive bayes, and random forest). Topics were only added to the record if all three methods agreed on the classification. The set size and test results using default tests from scikitlearn for each algorithm for each topic and subtopic for each of the five test runs can be found at: https://github.com/outbreak-info/topic_classifier/blob/main/results/in_depth_classifier_test.tsv

The efforts of our two volunteers suggested that non-experts were capable of thematically categorizing datasets, so we built a simple interface to allow citizen scientists to thematically classify the datasets that were available in our collection at that point in time. Each dataset was assigned up to 5 topics by at least three different citizen scientists to ensure quality of the results. Citizen scientists were asked to prioritize specific topic categories over broader ones. 90 citizen scientists recruited via either participation in the Mark2Cure project^38^ or a Scripps Research summer program participated in classifying 530 datasets pulled from Figshare and Zenodo, increasing the likelihood of quality submissions and decreasing the likelihood of abuse and false information. The citizen science curation site was originally hosted at https://curate.outbreak.info. The code for the site can be found at: https://github.com/outbreak-info/outbreak.info-resources/tree/master/citsciclassify. The citizen science classifications can be found at: https://github.com/outbreak-info/topic_classifier/blob/main/data/subtopics/curated_training_df.pickle. To evaluate the quality of the citizen scientist classifications, we first filtered classifications where at least ⅔ of 3-5 curators agreed on the topic category. We then compared the results of their classification with predictions by an out-the-box algorithm that was trained on LitCovid-classified abstracts. 186 of 530 classifications did not agree and were manually inspected; only about 10% of the categorization (54) was worse with citizen scientists over the predictions, and in many cases, the curators provided more precise categorization. Full details of the evaluation are available at https://github.com/gtsueng/curate_outbreak_data. These classifications have been incorporated into the appropriate datasets in our collection, and have been used to build our models for topic categorization. Basic in-group/out-group classification models were developed for each category using out-of-the-box logistic regression, multinomial naive bayes, and random forest algorithms available from SciKitLearn. The topic classifier can be found at https://github.com/outbreak-info/topic_classifier.

In addition to community curation of topic categorizations, we identified a citizen science effort, the COVID-19 Literature Surveillance Team (COVID-19 LST), that was evaluating the quality of COVID-19 related literature. The COVID-19 LST consists of medical students (many of which were in their third or fourth year), practitioners, and researchers who evaluate publications on COVID-19 based on the Oxford Levels of Evidence criteria and write Bottom Line, Up Front summaries^24^. With their permission, we integrated their outputs (daily reports/summaries, and Levels of Evidence evaluations) into our collection. Although the project has since ended, the valuable work by this team was integrated without further evaluation due to their background/training.

We further integrated our publications by adding structured linkage metadata, connecting preprints and their peer-reviewed versions. We performed separate Jaccard’s similarity calculations on the title/text and authors for preprint versus LitCovid Publications. We identified thresholds with high precision, low sensitivity and binned the matches into (expected match vs needs review). We also leveraged NLM’s pilot preprint program to identify and incorporate additional matches. The code used for the preprint-matching and the .xlsx file detailing the semiautomated and manual inspection of a sample of 1,500 matches from the results can be found at https://github.com/outbreak-info/outbreak_preprint_matcher. Briefly, a subsample of 1,500 matches were inspected and confirmed to match via PubMed identifier/correction mapping (1,158), manual inspection of preprint records (290), and manual inspection of preprint and the corresponding PubMed record and figures (52). The inspection confirmed that our threshold cutoff for preprint matching ensured the inclusion of a limited number of the most accurate matches at the cost of many more potential, but lower quality matches. Expected matches were linked via the correction property in our schema.

### Case study on variant research

To identify research about variants, we used the keyword phrase variant OR lineage in the Research Library and within the R package outbreakinfo. For **Figure 4a**, resources were counted by @type (Publication, Dataset, ComputationalTool, ClinicalTrial, Protocol, Analysis). The number of resources was aggregated to the weekly level by the date of the latest update and normalized to all resources within the Library for that week, creating a proportion of the Library for that week (**Figure 4c**). For variant-specific queries, the WHO-designated name was combined with its Pango lineage plus all descendants, as specified by the Pango team in October 2022 (https://raw.githubusercontent.com/cov-lineages/lineages-website/master/data/lineages.yml). To decrease the likelihood of a spurious hit for the resource (for instance, a Publication mentioning Alpha in the description but focusing only on Omicron), we used fielded queries to only search by the name of the resource. For instance, for Gamma, the following query was used: `name:Gamma OR name:“P.1” OR name;“P. 1.2”`. Code to replicate the analysis and visualizations is available at https://github.com/outbreak-info/outbreak-resources-paper/blob/main/Figure%204%20-%20Variant%20analysis.R.

### Harmonization and integration of resources and genomics data

The integration of genomics data from GISAID is discussed elsewhere^21^. We built separate API endpoints for our resources (metadata resources API) and genomics (genomics data API) using the BioThings SDK^23^. Data is available via our API at api.outbreak.info and through our R package, as described in Gangavarapu et al.

## Acknowledgements

We would like to thank Jasmine Rah, Brennan Joseph Enright, Justin Doroshenko, Thamanna Nishath and the rest of the COVID-19 Literature Surveillance Team for their incredible work and allowing us to share their work. We would like to thank Tom Adams and Craig Lazarchick for their work in identifying metadata from various individual datasets and their extensive feedback. We thank Sue Andarmani for her suggestions and feedback on dataset categories. We thank all of the contributors found at (https://blog.outbreak.info/dataset-topic-category-contributors) for taking the time to categorize datasets. We thank David Valentine for sharing details about his netlify app as part of the Radx-Radical Data Coordination Center which is funded by NIH (U24LM013755).

## Data availability statement

All metadata harvested and harmonized in the outbreak.info Research Library is freely available through an API (http://api.outbreak.info/) and in an associated R package (https://outbreak-info.github.io/R-outbreak-info/).

## Code availability statement

All code used to generate the outbreak.info Research Library is freely available on GitHub (https://github.com/outbreak-info) under open-source licenses. This code includes:

- *outbreak.info web application:* the code powering the outbreak.info front-end (https://github.com/outbreak-info/outbreak.info).
- *outbreak.info R package:* R package to access all the genomics and epidemiology data and Research Library metadata compiled and standardized on outbreak.info (https://github.com/outbreak-info/R-outbreak-info)
- *api.outbreak.info:* Code to create the application programming interface (API) to access Research Library metadata and cases & deaths data, available at api.outbreak.info (https://github.com/outbreak-info/outbreak.api)
- *bioRxiv and medRxiv:* Harvester of bioRxiv and medRxiv pre-print publications (https://github.com/outbreak-info/biorxiv)
- *ClinicalTrials.gov:* Harvester of clinical trials from clinicaltrials.gov (https://github.com/outbreak-info/clinical_trials)
- *COVID-19 LST:* Harvester of COVID-19 LST level of evidence ratings (https://github.com/outbreak-info/covid19_LST_reports)
- *COVID-19 LST Annotations* (https://github.com/outbreak-info/covid19_LST_annotations)
- *COVID-19 LST Report Data* (https://github.com/outbreak-info/covid19_LST_report_data)
- *Data Discovery Engine:* Harvester for manually curated metadata from the Data Discovery Engine (https://github.com/biothings/discovery-app/blob/master/scripts/outbreak.py)
- *Figshare:* Harvester from Figshare COVID-19 (https://github.com/outbreak-info/covid_figshare)
- *Harvard Dataverse:* Harvester for COVID-19 collection of Harvard Dataverse (https://github.com/outbreak-info/dataverses)
- *Imperial College:* Harvester for analyses by Imperial College London (https://github.com/outbreak-info/covid_imperial_college)
- *LitCOVID:* LitCOVID publication harvester (https://github.com/outbreak-info/litcovid)
- *PDB:* Harvester of metadata for SARS-CoV-2 structures from the Protein Data Bank (https://github.com/outbreak-info/covid_pdb_datasets)
- protocols.io: Harvester of protocol metadata from protocols.io (https://github.com/outbreak-info/protocolsio)
- *WHO Clinical Trials:* Harvester of clinical trials from WHO ICTR (https://github.com/outbreak-info/covid_who_clinical_trials/blob/master/parser.py)
- Research Library Schemas: Reusable schemas for Publications, Datasets, ClinicalTrials, Protocols, and Analyses and associated data mappings (https://github.com/outbreak-info/outbreak.info-resources)
- *Research Library Tools:* Reusable tools for parsers (https://github.com/outbreak-info/outbreak_parser_tools)
- *Altmetric:* code to look up Altmetric ratings for outbreak.info resources (https://github.com/outbreak-info/covid_altmetrics)
- *Preprint matcher:* Code to match preprints to their peer-reviewed publications (https://github.com/outbreak-info/outbreak_preprint_matcher)
- *Topic Classifier:* Machine learning classification of categories within the Research Library (https://github.com/outbreak-info/topic_classifier)
- *Clinical Trials mappings:* Mapping logic used to classify clinical trial records using clinical trial-specific metadata (https://github.com/gtsueng/outbreak_CT_classifier)
- *Evaluation of Citizen Scientist Efforts:* https://github.com/gtsueng/curate_outbreak_data
- *Figures within this Manuscript:* Figures generated here, including for the case study https://github.com/outbreak-info/outbreak-resources-paper

