## Supplemental Figures for "Outbreak.info Research Library: A standardized, searchable platform to discover and explore COVID-19 resources"

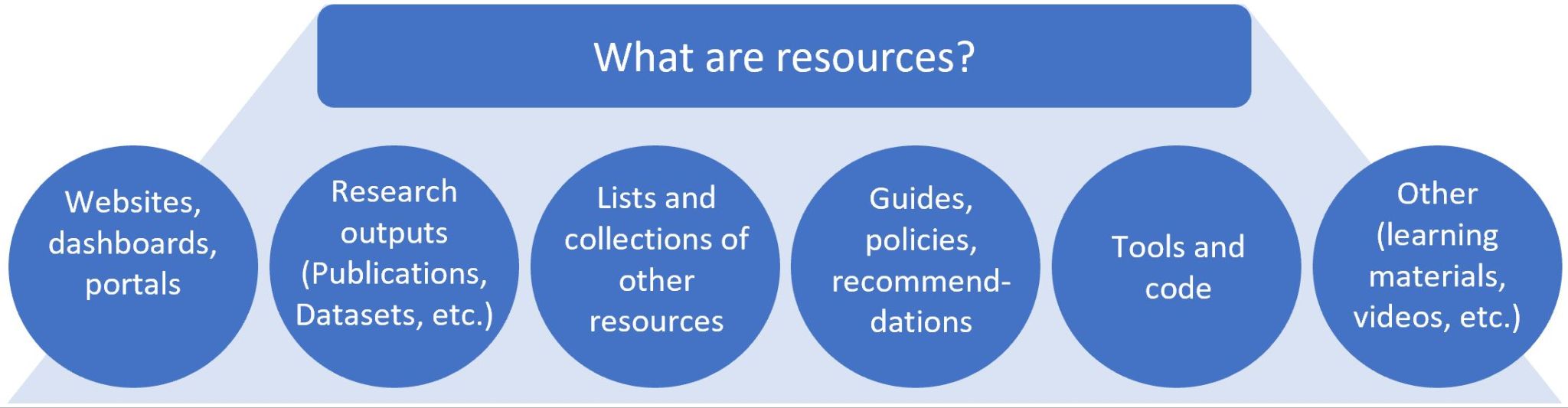

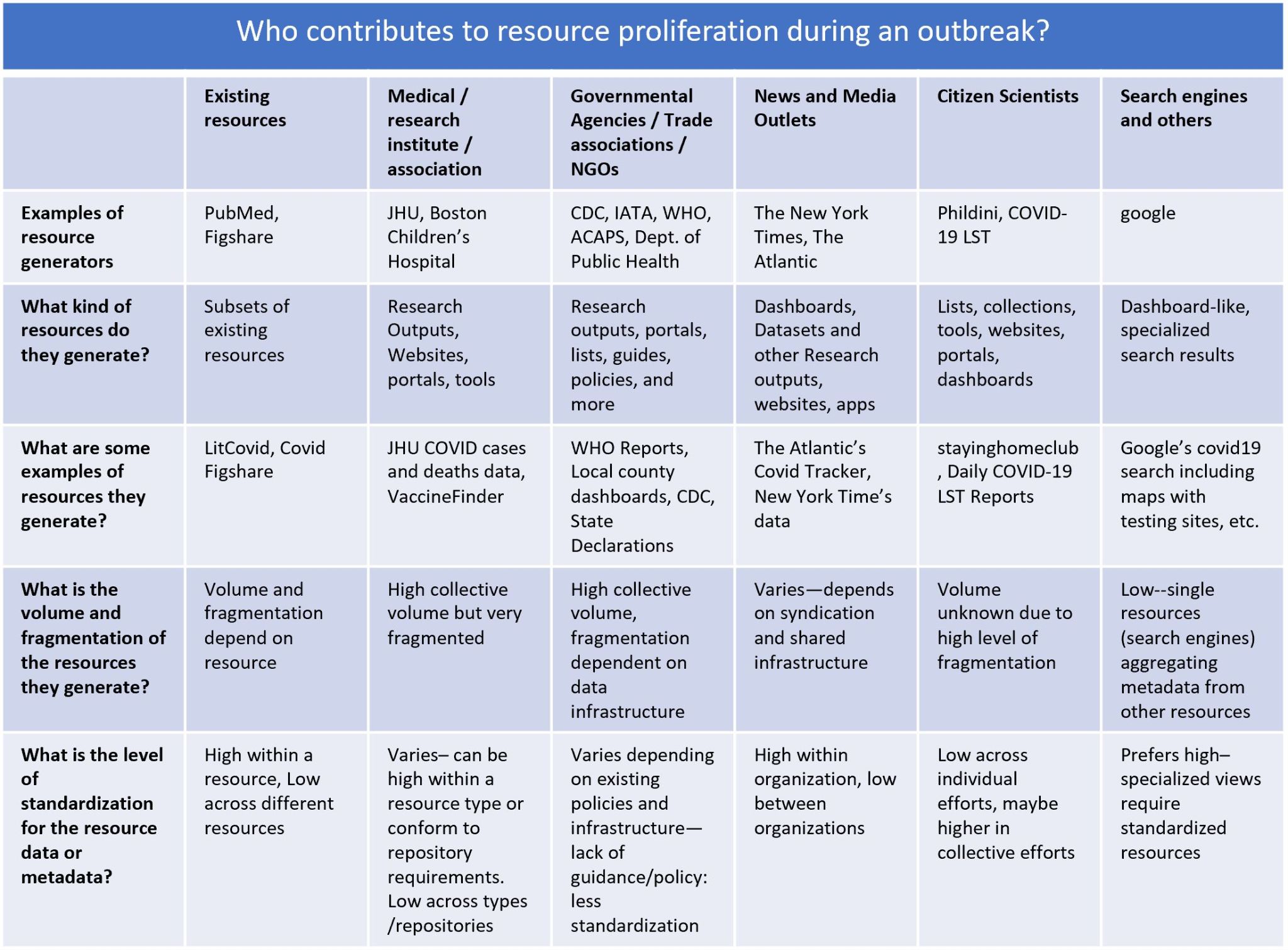

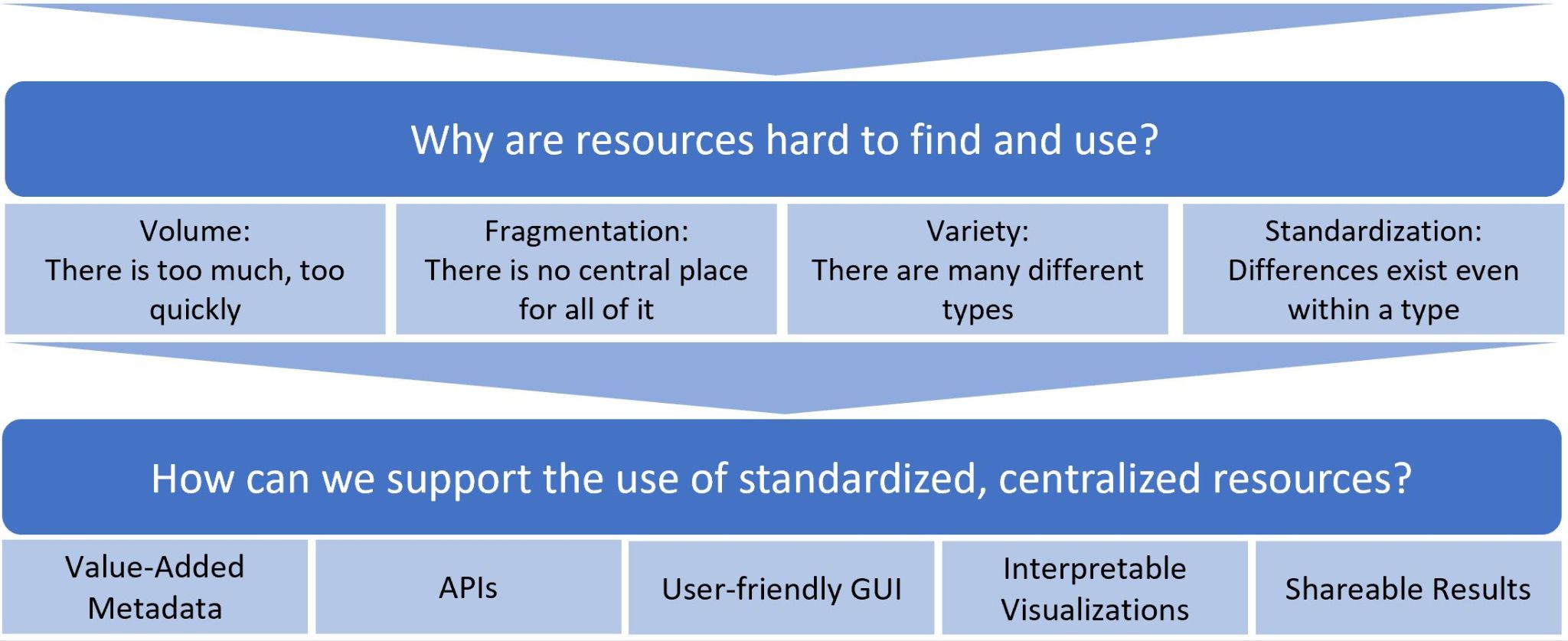

**Supplemental Figure 1.** What are resources, who contributes to the proliferation of resources, why are resources difficult to find and use, and how can we support their use?

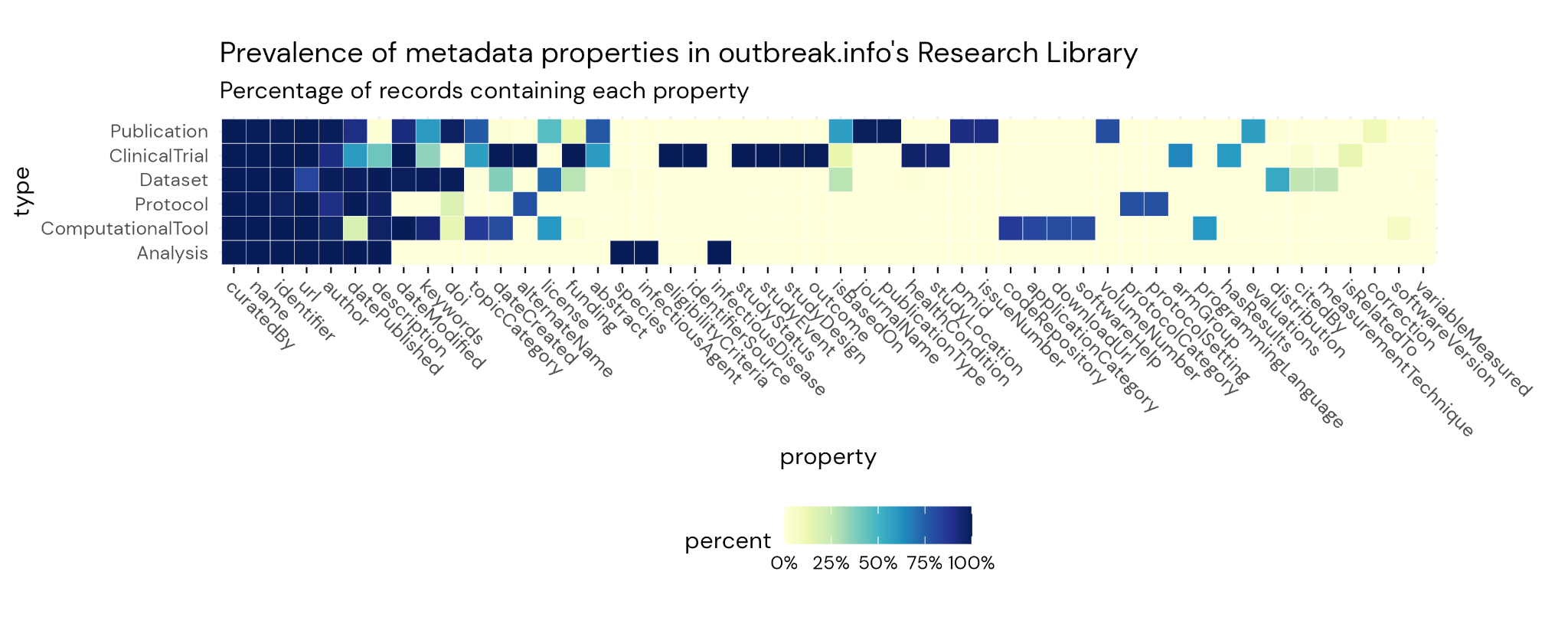

**Supplemental Figure 2**. Prevalence of each primary metadata field in our schemas by resource type. Raw data is provided in **Supplemental Table 1**.

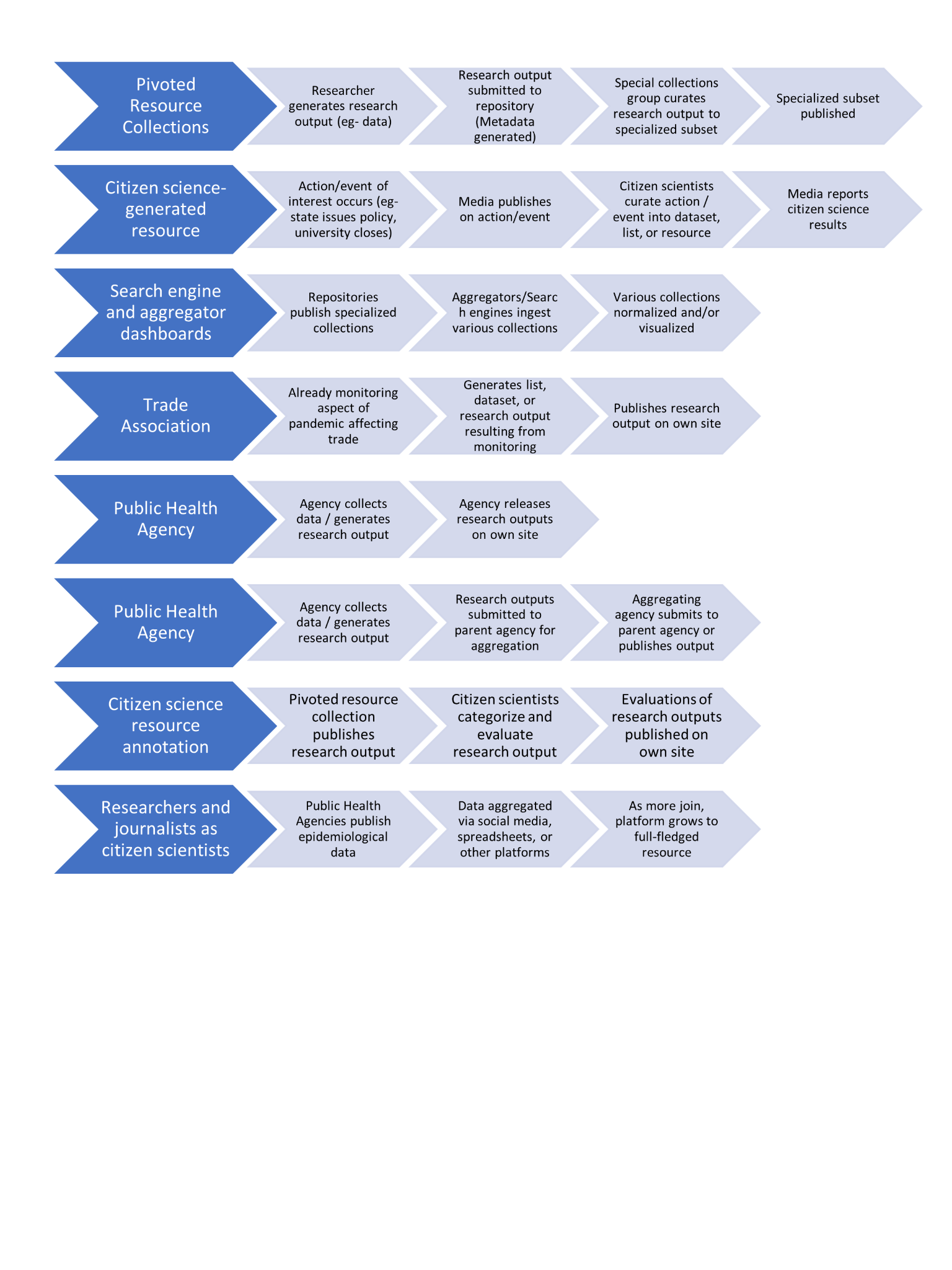

**Supplemental Figure 3**. Example resource-generation pipelines observed within the first month of the pandemic, resulting in a highly-fragmented, non-standardized resource landscape that is difficult to search, centralize and leverage.

### Usability Studies and Protocols

**A. Usability Study Results**

| Date | n | Task Number | SCR | EFR | TPT | EOU | LTR |
| --- | --- | --- | --- | --- | --- | --- | --- |
| Nov 23-25, 2020 | 5 | 1 | 100% | 60% | ~1 min | 96% | Highly likely |
|  |  | 2 | 100% | 0% | ~1 min | 100% | Highly likely |
| Mar 4 - Apr 2 , 2021 | 5 | 3 | 100% | 100% | 40 sec | 100% |  |

SCR- Average Successful Completion Rate for the task, EFR - Average Error-Free Rate, TPT- Average Time Per Task, EoU- Average Ease of Use Rating, LTR - Average Likelihood To Return Rate

**B. Usability Study Protocol**

*Audience*: Biomedical researchers

*Purpose*: Test the ease by which biomedical researchers could use outbreak.info to access or visualize data and find up-to-date publications.

*Format*: Moderated usability study conducted via Zoom. Users were asked to perform one or more of the following tasks:

1. Please use the Research Library to find new datasets related to the COVID-19 reproduction number in the United States.
2. Please use the Research Library to find publications related to cytokine response storm in COVID-19 patients.
3. Please find all publications related to B.1.1.7.

*Post-test questions asked:*

- Overall, please rate how easy or difficult it is to use outbreak.info on a scale of 1-5, where 1 is very difficult and 5 is very easy.
- How likely are you to continue using outbreak.info to regularly access data or resources, on a scale of 1-5 where 1 is very unlikely and 5 is very likely?
- What do you like most about the outbreak.info app?
- What would you improve about the app? Or what would you add to the app?
- How would you compare outbreak.info to other sites you’ve used to access data & resources?

*Metrics/Measurements recorded*:

- Ease and satisfaction about each task (5-point Likert scale),
- Time on each task,
- Number of successful task completions and errors,
- Overall ease and satisfaction (5-point Likert scale),
- Likelihood to use (5-point Likert scale),
- Suggestions for improvement (likes, dislikes, recommendations),
- Error-free rate,
- And noted observations about the users’ process.

**C. Changes ensure usability of Research Library**

| Usability Study | Changes Implemented |
| --- | --- |
| Nov 23-25, 2020 | - Redesigned homepage to present outbreak.info’s main features as three actionable categories near the top - Added an introductory video - Revised how other pages are linked from the homepage - Collapsed many of the site’s details into expandable cards - Renamed “Resources” to “Research Library.” |
| Mar 4 - Apr 2 , 2021 | - No changes were made to the Research Library |

**Supplemental Figure 4.** Usability studies for iterative design improvements of the Outbreak.info Research Library. **A**. Remote moderated usability test of outbreak.info was conducted between November 23-25, 2020 and between March 4, 2021 and April 2, 2021 over Zoom. Five biomedical researchers participated in each round of tests which consisted of asking users to perform the tasks (and recording metrics) in accordance with protocol (**B**). There was a successful completion rate of 100% for both tasks and zero critical errors, but some non-critical errors resulted in an error-free rate of 60% for the first task and 0% for the second task. The average amount of time spent on each task was around 1 minute. **C.** Improvements to the site and Research Library were made with accordance to the results of the usability studies in November of 2020, but follow-up usability studies in March-April of 2021, indicated no additional changes were needed in order to improve the user performance on the tested tasks.

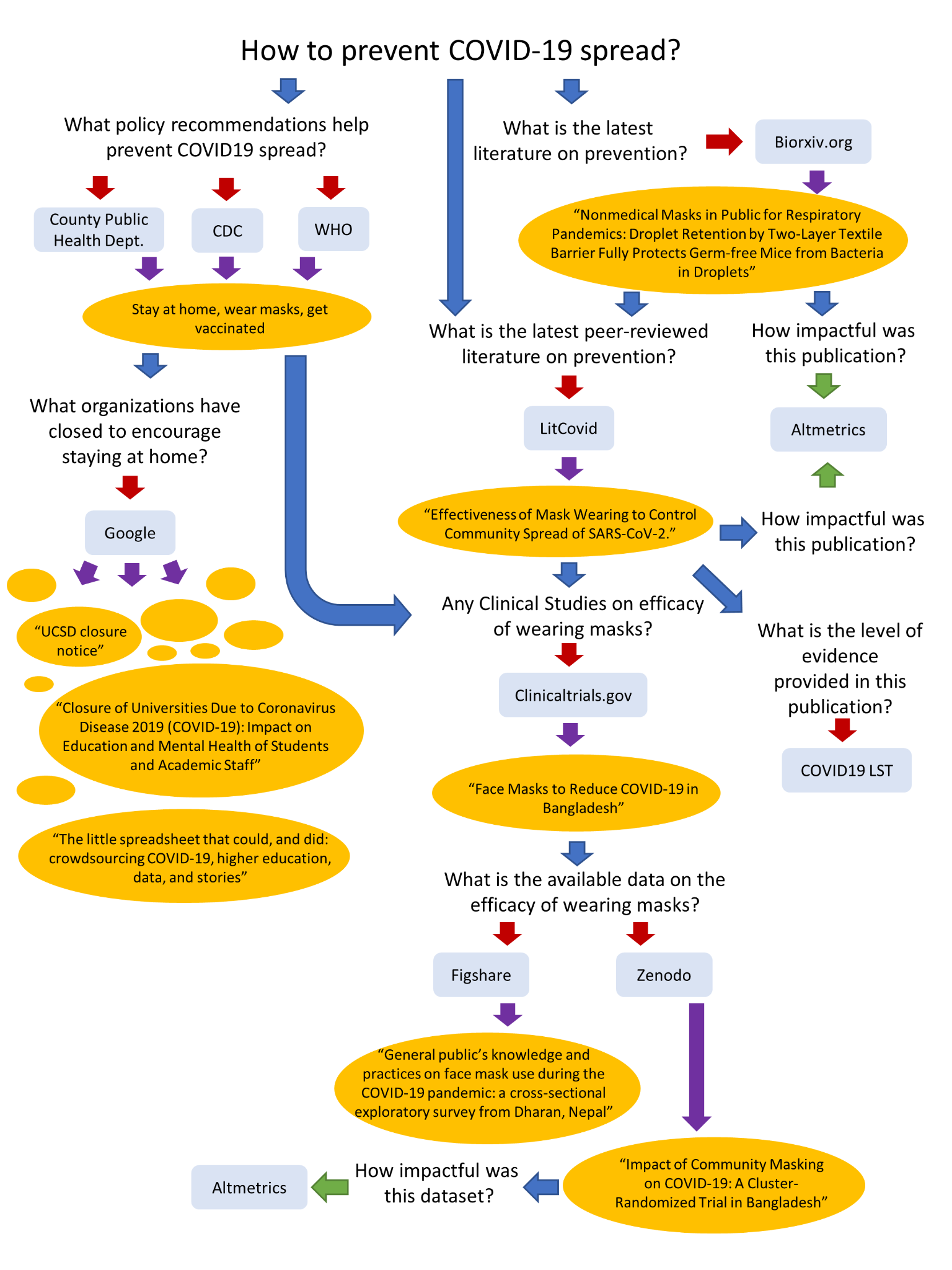

**Supplemental Figure 5.** Example resource exploration pathway for addressing the question, “How to prevent COVID-19 spread” early on in the pandemic. The question leads to an iterative process of refining the question (unbound text), searching a resource/website (blue rounded box), finding a result of interest (orange oval). Blue arrows indicate a question refinement step. Purple arrows indicate a result review step (i.e.- identifying a potential result of interest). Red arrows indicate a manual search and filter step. Green arrows indicate linked search (i.e.- manual input to a resource not needed for result).

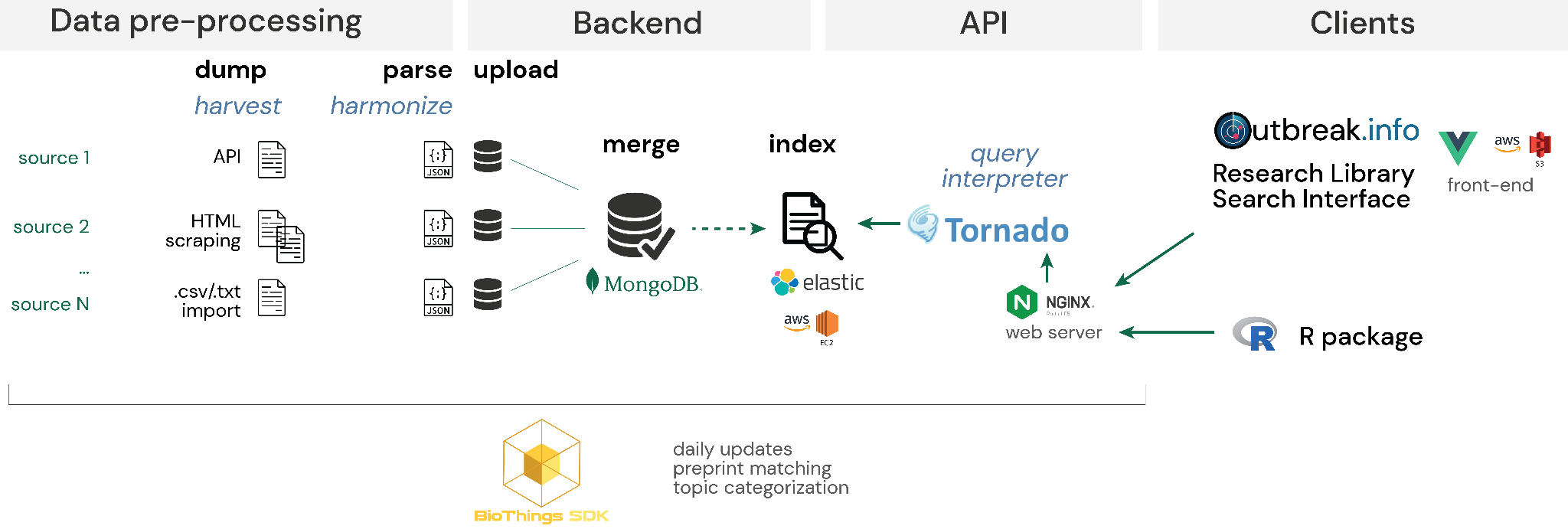
**Supplemental Figure 6**. Systems architecture for the outbreak.info Research Library metadata infrastructure. Individual data sources are harvested either through API-based calls, reading a tabular file or scraping-based HTML crawling, harmonized to our schemas through custom parser scripts, and uploaded, merged, and indexed to an Elasticsearch index using BioThings SDK. Since each source is harvested and harmonized independently, errors in a given data source do not affect updates in the other sources or the overall status of the API. Queries from the Research Library front-end or the R package are routed through a Tornado-based interpreter which accesses the underlying Elasticsearch index to return results from the API call.
